# Manual stabilization reveals a transient role for balance control during locomotor adaptation

**DOI:** 10.1101/647453

**Authors:** Sungwoo Park, James M. Finley

**Affiliations:** Division of Biokinesiology and Physical Therapy, University of Southern California, Los Angeles, CA, 90033; Neuroscience Graduate Program, University of Southern California, Los Angeles, CA, 90089

**Keywords:** Gait, Adaptation, Motor learning, Balance, Stability

## Abstract

A fundamental feature of human locomotor control is the need to adapt our walking pattern in response to changes in the environment. For example, when people walk on a split-belt treadmill which has belts that move at different speeds, they adapt to the asymmetric speed constraints by reducing their spatiotemporal asymmetry. Here, we aim to understand the role of stability as a potential factor driving this adaptation process. We recruited 24 healthy, young adults to adapt to walking on a split-belt treadmill while either holding on to a handrail or walking with free arm swing. We measured whole-body angular momentum and step length asymmetry as measures of dynamic balance and spatiotemporal asymmetry, respectively. To understand how changes in intersegmental coordination influenced measures of dynamic balance, we also measured segmental angular momenta and the coefficient of limb cancellation. When participants were initially exposed to the asymmetry in belt speeds, we observed an increase in whole-body angular momentum that was due to both an increase in the momentum of individual limb segments and a reduction in limb cancellation. Holding on to a handrail reduced the perturbation to asymmetry during the early phase of adaptation and resulted in a smaller after-effect during post-adaptation. In addition, the stabilization provided by holding on to a handrail led to reductions in the coupling between angular momentum and asymmetry. These results suggest that regulation of dynamic balance is most important during the initial, transient phase of adaptation to walking on a split-belt treadmill.

**Summary Statement:** Regulation of balance exhibits a transient effect on adaptation to imposed asymmetries during bipedal walking. External stabilization attenuates initial deviations in spatiotemporal asymmetry but has no effect on subsequent adaptation.

## Introduction

The ability to successfully adapt our movements in variable environments is a hallmark of skilled motor control. This process of motor adaptation requires the nervous system to form new calibrations relating motor actions to a set of desired objectives. Locomotor adaptation has most frequently been explored in the context of adaptation to walking on a split-belt treadmill which has two independent belts capable of moving at different speeds (Darter et al., 2018; Dietz et al., 1994; Reisman et al., 2005; Sánchez et al., 2017; Selgrade et al., 2017). The adaptation process is characterized by a gradual reduction in step length asymmetry that is accompanied by a concomitant reduction in metabolic cost (Finley et al., 2013; Selgrade et al., 2017) and positive mechanical work. Thus, the final behavior that people adopt during adaptation is more economical than the strategy used when they are initially exposed to the asymmetry in belt speeds. However, since balance is one of the primary challenges of bipedal locomotion, a complete understanding of locomotor adaptation requires an analysis of the causal link between perturbations in speed, balance control, and spatiotemporal asymmetry.

Recent work has shown that when people adapt to walking on a split-belt treadmill, they increase the fore-aft margins of stability when stepping on the fast belt (Buurke et al., 2018; Darter et al., 2018; Park and Finley, 2017). These margins represent the distance between the edge of base of support and the extrapolated center of mass (XCoM, (Hof et al., 2005)) and increasing fore-aft margins of stability provides more time for the CoM to reach to the edge of base of support before the next step is required (Slobounov et al., 1998). Although these studies show that control of foot placement relative to the CoM is associated with changes in spatiotemporal asymmetry during walking on a split-belt treadmill, margins of stability do not capture the full dynamics of the body as they relate to potential fall risk.

Measures of whole-body angular momentum (WBAM) about the body’s CoM have been used to capture the body’s rotational dynamics during walking (Elftman, 1939; Herr and Popovic, 2008). The peak-to-peak range of WBAM over a stride is typically regulated to stay within a small range during normal walking due to the cancelation of momentum between contralateral limb segments. In addition, when people are exposed to an unexpected perturbation during walking, there is an increase in WBAM (Liu et al., 2018; Martelli et al., 2013) which may result from an increase in whole-body rotation or a reduction in contralateral cancellation of segmental angular momenta. Although dynamic balance appears to be regulated by controlling WBAM during normal walking, how WBAM is regulated during split-belt adaptation has yet to be determined. If a desire to maintain balance is one of the factors that drive adaptation, measures of WBAM should be correlated with measures of spatiotemporal asymmetry. For example, WBAM may scale with the magnitude of asymmetry due to an increase in whole-body rotation or a reduction in cancellation of contralateral segmental momenta.

Establishing the relationship between asymmetry and measures of stability may provide evidence for the role of balance regulation during adaptation but identifying casual associations can be probed more directly through the provision of external support to reduce the need for active control of stability. When people use an external source of stability such as a handrail or cane during walking, they reduce step length variability and reduce the excursion of the CoM within the base of support (Bateni and Maki, 2005; Dickstein and Laufer, 2004). Even light touch of a handrail leads to reductions in fore-aft and mediolateral body sway during standing (Dickstein and Laufer, 2004; Jeka, 1997; Jeka and Lackner, 1994). If asymmetry is a correlate of instability, provision of an external source of stability should be accompanied by a reduction in spatiotemporal asymmetry during initial exposure to the split-belt perturbation. Similarly, when the belts are driven to move at the same speed during post-adaptation, the initial aftereffect should be smaller than that measured when participants are not provided with a source of support.

Here, we examined changes in dynamic balance during split-belt adaptation both with and without an external source of stability. We hypothesized that people would increase WBAM during early adaptation compared to the baseline due to increases in whole-body rotation and reductions in the degree of cancellation of contralateral limb rotations due to asymmetries in inter-limb coordination. We also hypothesized that over the course of adaptation, WBAM would decrease as people reduce step length asymmetry. When provided with an external source of stability through use of a handrail, we hypothesized that people would reduce step length asymmetry during early adaptation and early post-adaptation and reduce WBAM during early adaptation and early post-adaptation. Together these experiments will help us better understand the role of stability in driving changes in spatiotemporal asymmetry during locomotor adaptation.

## Methods

### Participants

A total of 24 healthy young individuals (age 27±2.3, 6 female) were recruited for this study. Participants were randomly assigned to a group that walked on the treadmill without holding on to a handrail (Hands Off group, n = 12) or a group that walked while holding on to a handrail in front on the treadmill (Hands On group, n = 12). We based our sample size calculation on a set of pilot data (n = 10) comparing WBAM during tied-belt walking and during the early phase of adaptation on a split-belt treadmill. The effect size and standard deviation for this pilot data were 0.15 and 0.05, respectively. The desired power and alpha level were set to 80% and 0.05. All experimental procedures were approved by the University of Southern California Institutional Review Board and each participant provided written, informed consent before testing began. All aspects of the study conformed to the principles described in the Declaration of Helsinki. All participants provided written informed consent before testing.

### Experimental protocol

Participants initially walked on the treadmill at three different speeds for 5, 2.5, and 2.5 minutes at 1.0, 0.5, and 1.5 ms^-1^, respectively (Fig. 1). During adaptation, the belts moved at 0.5 and 1.5 ms^-1^, respectively and the fast side was randomized across participants. These belt speeds were chosen to be consistent with previous work (Finley et al., 2013, 2014, 2015; Malone et al., 2012). Participants walked for 15 minutes with the belts split and were instructed not to look down at the belts. They then walked for 10 minutes with the belts tied at 1.0 ms^-1^ during post-adaptation to allow us to evaluate the persistence of adaptive changes in their walking pattern when the perturbation was removed. The Hands On group held on to a handrail throughout the experiment for all time phases, while the Hands Off group did not hold on to the handrail at all. Participants in the Hands On group were instructed not to lean on the handrail but to only rest their hands on it. There were no rest periods between trials.

**Figure 1.**
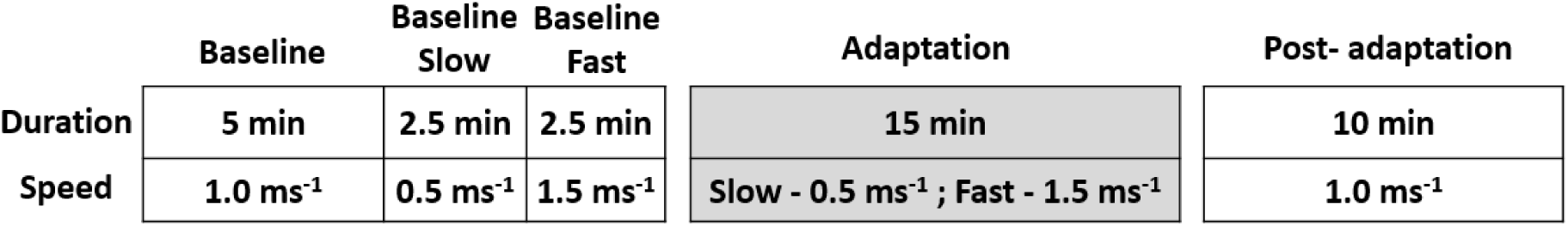
Experimental protocol. Participants completed three baseline trials at walking speeds of 1.0, 0.5, and 1.5 ms^-1^. During adaptation, participants walked with the belts moving at a 3:1 speed ratio (0.5 and 1.5 ms^-1^) for 15 minutes. Participants then walked for 10 minutes with the belts tied at 1.0 ms^-1^ during post-adaptation.

### Data acquisition

Segmental kinematics during treadmill walking were recorded using a full body, passive marker set (Liu et al., 2018; Song et al., 2012). Retroreflective markers were placed on the following bony landmarks: anterior, posterior, and lateral cranium, acromion processes, anterior and posterior shoulders, greater tubercles of humerus, medial and lateral humeral epicondyles, radial styloid process, ulnar head, 7th cervical vertebrae, sternoclavicular notch, iliac crests, anterior superior iliac spines, posterior superior iliac spines, L5-S1 joint, medial and lateral femoral epicondyles, medial and lateral malleoli, and first and fifth metatarsal heads. In addition, sets of cluster markers were placed over the upper arms, lower arms, thighs, shanks, and heels. Marker kinematics were recorded using a 10-camera Qualisys Oqus motion capture system (Qualisys AB, Goteborg, Sweden).

### Data processing

Markers were labeled using Qualisys Track Manager and then exported to Visual 3D (C-Motion, Rockville, MD, USA) to construct a full body model. Kinematic variables associated with spatiotemporal gait asymmetry were calculated using 3D marker positions (Finley et al., 2013; Reisman et al., 2005). Step lengths were calculated by measuring the distance between heel markers at foot heel strike. Step length asymmetry (SLA) was computed as the normalized difference in step lengths based on Eqn 1. Here, *SL*_*fast*_ is the step length at foot strike when the leading leg is on the faster belt and *SL*_*slow*_ is the step length at foot strike when the leading leg is on the slower belt.

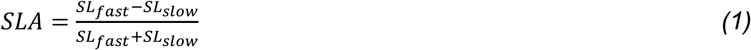

We calculated WBAM about the CoM in the sagittal plane as a measure of dynamic balance during walking. WBAM 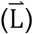 is the sum of the angular momenta of individual limb segments with respect to the whole-body CoM (Herr and Popovic, 2008), and nondimensionalized by body mass (M), the CoM height (H), and the term 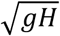, where *g* is the gravitational acceleration (Vistamehr et al., 2016). The term 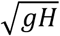 has units of ms^-1^ and is independent of walking speed.

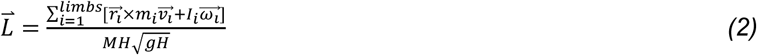

The angular momentum of the individual limb segments in the sagittal plane was calculated as the sum of the angular momentum of each segment about a mediolateral axis projecting through the body’s CoM (*r*_*i*_: position of the segmental CoM, m_i_: segment mass,*v*_*i*_: linear velocity of the segment relative to body’s CoM) and the angular momentum of each segment about its own CoM (*I*_*i*_: segmental moment of inertia, *ω*_*i*_: segmental angular velocity).

### Data analysis

We calculated the integrated angular momentum (*ΣL*) during each step cycle by computing the area under the angular momentum curve (Fig. 2). *ΣL* captures the cumulative effects of whole-body rotation and interlimb cancellation during walking. We analyzed *ΣL* over each step cycle throughout each phase of the experiment. We took the first five strides of adaptation and post-adaptation to represent the early phase (Early Adaptation and Early Post-Adaptation) and the last five strides to represent the late phase (Late Adaptation and Late Post-Adaptation). Thus, there were seven time phases including Baseline, Baseline Slow, Baseline Fast, Early Adaptation, Late Adaptation, Early Post-Adaptation, and Late Post-Adaptation for the average data.

**Figure 2.**
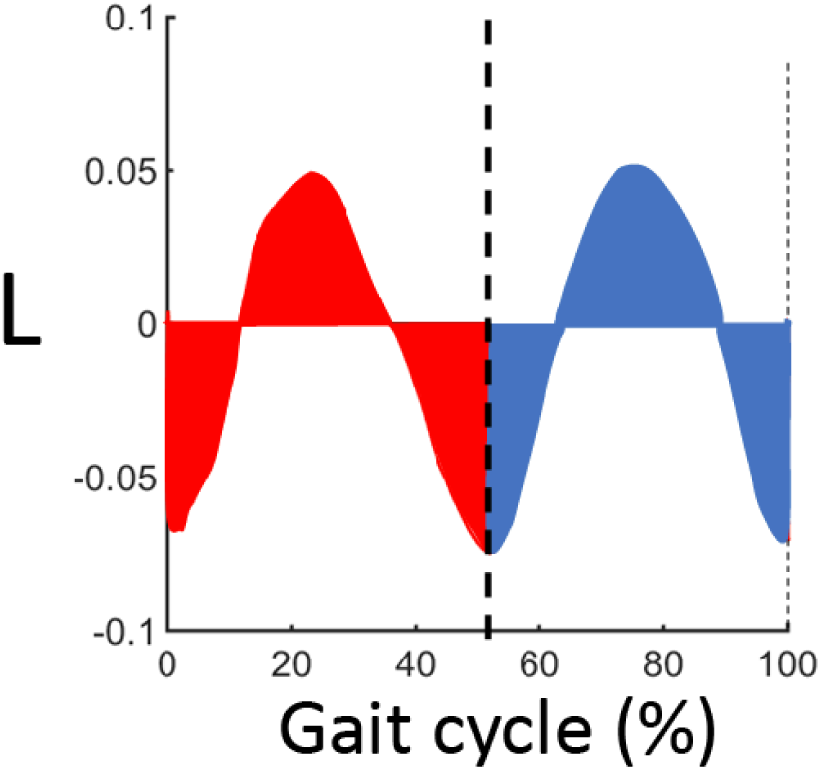
Whole-body angular momentum in the sagittal plane (L) over a gait cycle. Integrated angular momentum for each step cycle (red and blue, respectively) was calculated by computing the area under the curve during each step.

WBAM is regulated in part by the fact that contralateral limbs move in anti-phase such that analogous limb segments cancel each other’s angular momenta (Herr and Popovic, 2008). Here, we quantified this phenomenon using the coefficient of cancellation, *k*, over a gait cycle (Bennett et al., 2010).

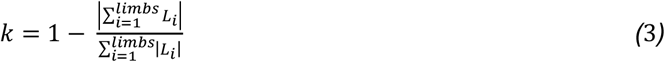

*L*_*i*_ is the angular momentum in the sagittal plane of the *i*th body segment. If all segments cancelled one another perfectly, *k* would be 1. Conversely, if there was no interlimb cancellation, *k* would be 0 since the numerator and denominator would be equal. For the statistical analysis, we computed the mean value of the coefficient of cancellation over each stride.

### Statistical analysis

All statistical analyses were performed in Matlab version R2018a (Mathworks, Natick, MA, USA). We tested the hypotheses that the magnitude of the integrated whole-body angular momentum (WBAM) would be greater during Early Adaptation compared to Baseline and would each decrease over the course of adaptation. We also tested the hypothesis that walking while holding on to a handrail would decrease WBAM across all time periods, lead to a reduction in step length asymmetry, and reduce the coefficient of cancellation during Early Adaptation and Early Post-Adaptation. We tested these hypotheses using repeated measures ANOVAs to determine if there was a main effect of Phase (Baseline, Baseline Slow, Baseline Fast, Early Adaptation, Late Adaptation, Early Post-Adaptation, and Late Post-Adaptation), Group, or an interaction between Phase and Group on WBAM and SLA. We conducted post-hoc pairwise comparisons using Tukey’s honest significant difference criterion.

Lastly, we calculated Pearson’s correlation coefficients between SLA and measures of WBAM during adaptation and post-adaptation to determine if SLA and measures of WBAM were associated with each other and if this association differed across handrail conditions. Correlation coefficients were calculated for the first 50 strides during adaptation and post-adaptation since this was the timespan of the task during which SLA changed most, and yielded values of WBAM and SLA that were normally distributed. We used a two-way ANOVA to determine if the correlations between SLA and measures of WBAM differed across handrail conditions or time phases.

## Results

### Changes in step length asymmetry during split-belt adaptation

Over the course of adaptation, people gradually reduced the magnitude of SLA and they exhibited an after-effect and gradual decay in SLA when the belts were driven to move at the same speed again during post-adaptation (Fig. 3A). Participants in the Hands Off group had a large initial asymmetry during Early Adaptation, but reduced this asymmetry during Late Adaptation relative to Early Adaptation (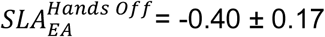 versus 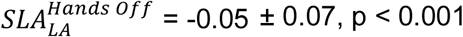). During Early Post-Adaptation, participants exhibited an aftereffect consisting of an increase in SLA in the opposite direction of what was observed during Early Adaptation 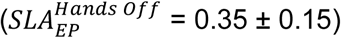. They restored SLA to Baseline levels by Late Post-Adaptation 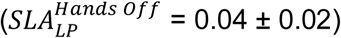. Holding on to the handrail reduced the magnitude of SLA during Early Adaptation (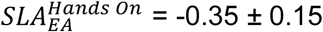 versus 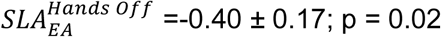) and Early Post-Adaptation (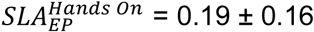 versus 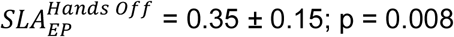).

**Figure 3:**
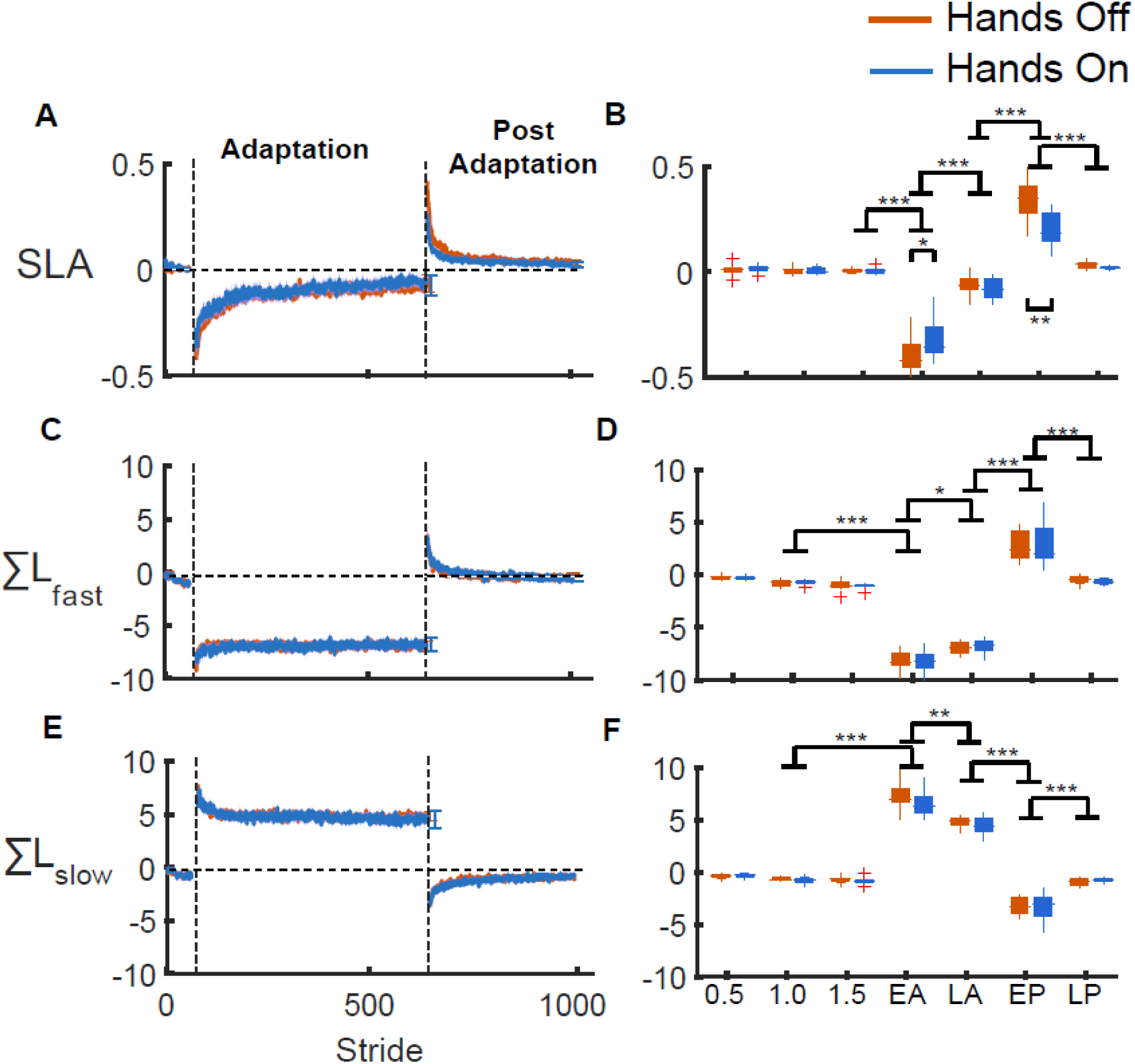
Stride-by-stride and average values of step length asymmetry and integrated WBAM. A and B. Step length asymmetry, C and D. integrated WBAM over a step cycle for the limb that was on the fast belt during adaptation, E and F. integrated WBAM over a step cycle for the limb that was on the slow belt during adaptation. For time series data (left column), the solid lines represent the median across participants and the shaded area represents the standard deviation. Red: Hands off group. Blue: Hands on group. *p < 0.05; **p<0.01; ***p<0.001).

### Regulation of WBAM throughout the gait cycle

WBAM varied throughout the gait cycle and captured the cyclic pattern of forward falling and recovery that characterizes bipedal walking (Fig. 4). During baseline walking, the trajectory of WBAM during each half of the gait cycle was symmetric around zero, indicating equal forward and backward rotation within each step cycle. Throughout adaptation, the WBAM trajectory during the stance phase on the fast belt remained negative which indicates that participants tended to fall forward throughout the fast step cycle (Fig. 4D and E). They recovered from this forward rotation by increasing backward rotation (positive WBAM) during the step on the slow belt. During Early Post-Adaptation, WBAM was more positive during the first half of the gait cycle than the second (Fig. 4F). This indicated that people tended to fall forward more during the step on the former slow belt than during the step cycle on the former fast belt. They then returned to the trajectory observed during baseline walking at the average belt speeds by Late Post-Adaptation (Fig. 4G).

**Figure 4:**
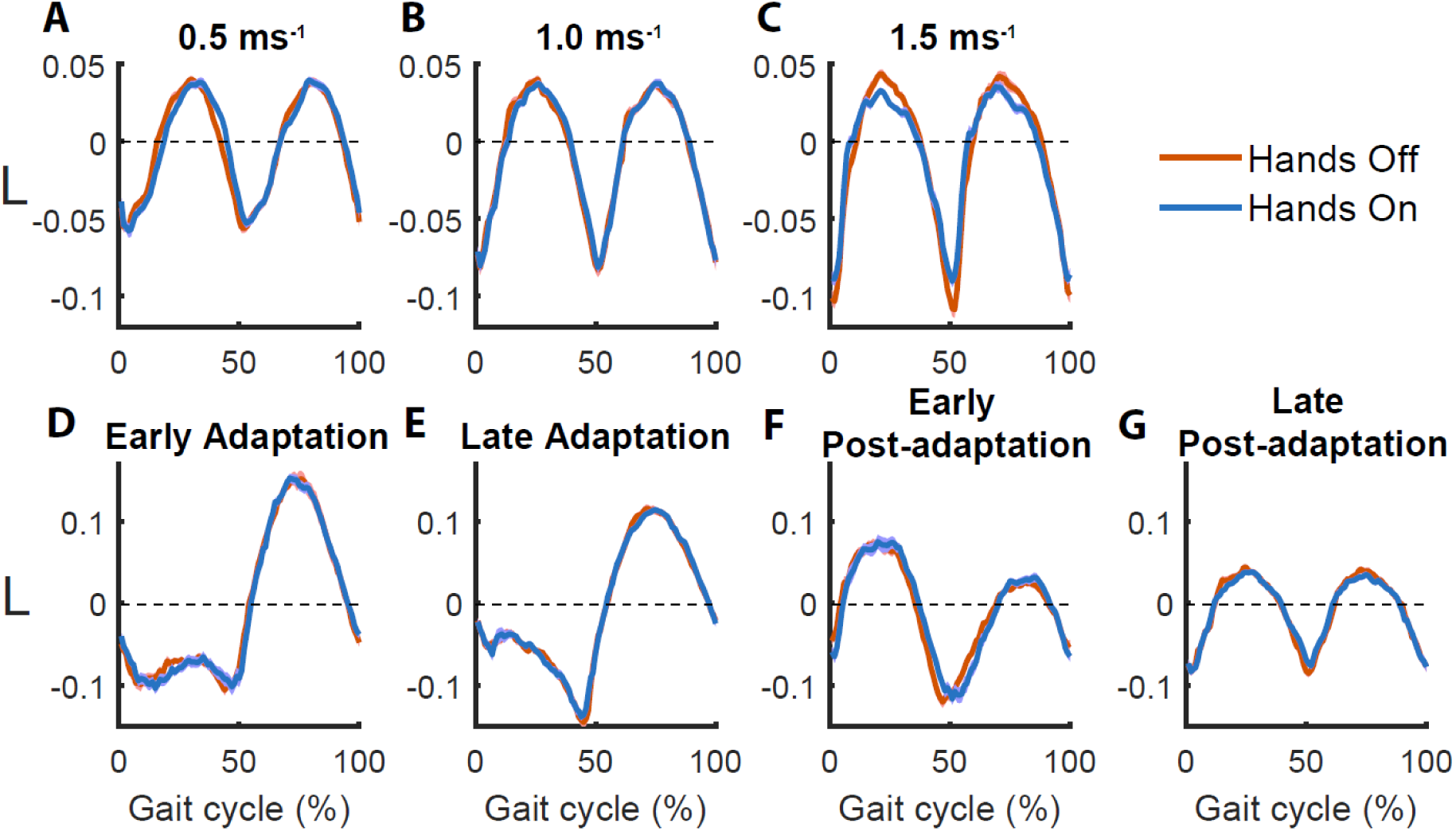
Whole-body angular momentum over the gait cycle for A. 0.5 ms^-1^, B. 1.0 ms^-1^, C. 1.5 ms^-1^, D. Early Adaptation, E. Late Adaptation, F. Early Post-Adaptation, and G. Late Post-Adaptation. During Early and Late Adaptation, gait cycles start with foot strike on the fast belt such that the first half of the gait cycle corresponds to stance phase on the fast belt and the second half corresponds to stance phase on the slow belt. During Post-adaptation, the gait cycle begins with the step on the side of the treadmill that was faster during Adaptation. The solid lines represent the median across participants and the shaded area around each curve represent the standard deviation across participants. Red: Hands Off. Blue: Hands On.

### Changes in integrated WBAM during adaptation

We quantified the changes in whole-body dynamics by measuring the integrated WBAM (*ΣL*) during steps on the fast and slow belts, respectively. During Early Adaptation, *ΣL* became more negative compared to Baseline when stepping on the fast belt (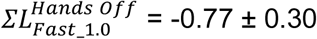 versus 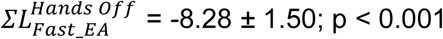). In contrast, *ΣL* became more positive compared to Baseline during steps on the slow belt (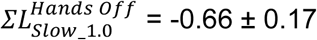 versus 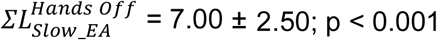). By the end of adaptation, there was a significant reduction in the magnitude of *ΣL* for both the fast (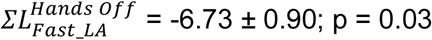) and slow steps (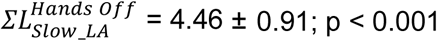). During Early Post-Adaptation, the direction of *ΣL* was opposite of that observed during Early Adaptation. *ΣL* during the step on the former fast belt became positive relative to Late Adaptation 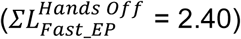 while *ΣL* during step on the former slow belt became more negative 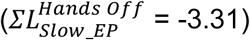. Ultimately, *ΣL* for both steps returned to Baseline level by the end of the post-adaptation period. Contrary to our hypothesis, holding on to the handrail did not affect *ΣL* during any phase of the experiment.

### Changes in limb angular momenta and segmental cancellation during adaptation

Although we expected that the provision of an external source of stability would reduce whole-body angular momentum during adaptation, this is not what we observed. To better understand the effects of manual stabilization on whole-body momentum, we examined differences in segmental momentum and interlimb cancellation between groups. This analysis revealed that although the angular momenta of the head and trunk systematically changed with belt speed asymmetry they did not differ across handrail conditions (Fig. 5). There was a significant effect of Phase on the integrated momentum of the head and trunk during both the fast and slow steps (F = 5.65, p = 0.02 and F = 9.31, p = 0.005, respectively), but there was no effect of Group during either the fast or slow step (F = 0.56, p = 0.46 and F = 0.81, p = 0.38, respectively), nor was there an interaction between Phase and Group when stepping on the fast or slow belts (F = 0.30, p = 0.59 and F = 0.33, p = 0.57, respectively). This indicates that holding on the handrail had no effect on sagittal plane momentum of the head and trunk. Similarly, while there was a significant effect of Phase on the integrated angular momentum of the legs during steps on the fast and slow belts (Fig. 5, F = 26.25, p < 0.001 and F = 20.67, p < 0.001, respectively), there was no effect of Group on momentum of the legs during steps on the fast or slow belt (F = 1.11, p = 0.30 and F = 1.87, p = 0.19, respectively), nor was there a significant interaction between Phase and Group for steps on the fast or slow belts (F = 1.18, p = 0.29 and F = 1.66, p = 0.21, respectively).

**Figure 5:**
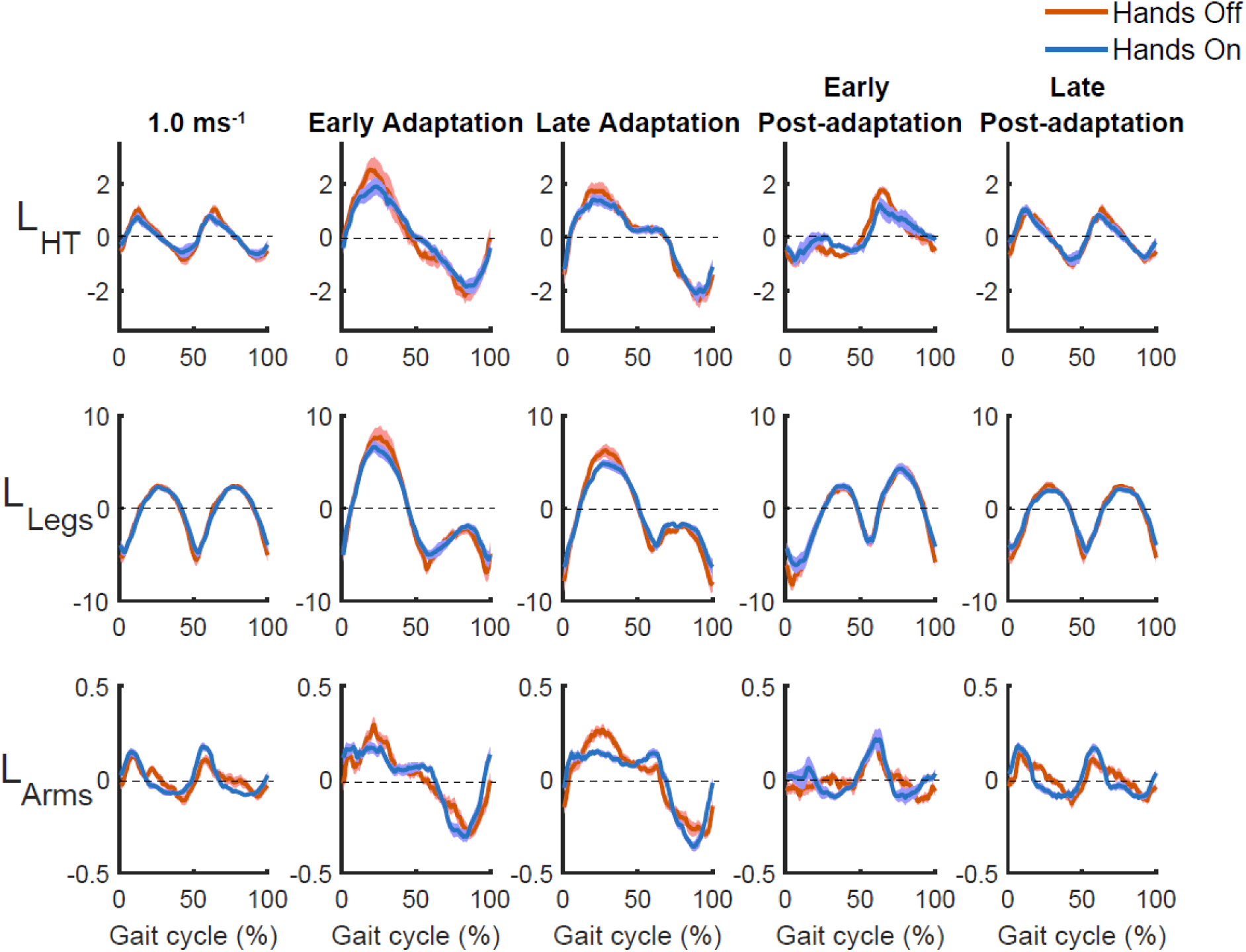
Angular momentum of HT (head and trunk), Legs (left and right thigh, shank, and foot), Arms (left and right upper-arm and fore-arm) over the gait cycle during Baseline (1.0 ms^-1^), Early Adaptation, Late Adaptation, Early Post-Adaptation, and Late Post-Adaptation. During Early and Late Adaptation, gait cycles start with foot strike on the slow belt such that the first half of the gait cycle corresponds to stance phase on the slow belt and the second half corresponds to stance phase on the fast belt. During Post-adaptation, the gait cycle begins with the step on the side of the treadmill that was slower during Adaptation. The solid lines represent the median across participants and the shaded area around each curve represent the standard deviation across participants. Red: Hands Off. Blue: Hands On.

We observed similar patterns for segmental cancellation (Fig. 6). There was a main effect of Phase on limb cancellation (F = 2118.6, p < 0.001), but there was no effect of Group (F = 1.03, p = 0.32) and no interaction between Phase and Group (F = 0.78, p = 0.39). The mean value of *k* over the gait cycle decreased during Early Adaptation compared to Baseline (*k*_1.0_ = 0.76 ± 0.03 versus *k*_*EA*_= 0.56 ± 0.03, p < 0.001), but *k* increased over the course of adaptation (*k*_*EA*_= 0.56 ± 0.03 versus *k*_*LA*_ = 0.62 ± 0.03, p = 0.004) likely as a result of changes in interlimb phasing. However, during Late Adaptation *k* was still smaller than during Baseline (p < 0.001) and this likely reflected the asymmetry in belt speeds. When the belts were set to move at the same speed again during post-adaptation, interlimb cancellation increased relative to Late Adaptation (*k*_*EP*_ = 0.72 ± 0.02, p < 0.001) and gradually returned to baseline values. These results indicate that holding on a handrail had no effects on whole-body angular momentum, segmental momentum, or segmental cancellation.

**Figure 6:**
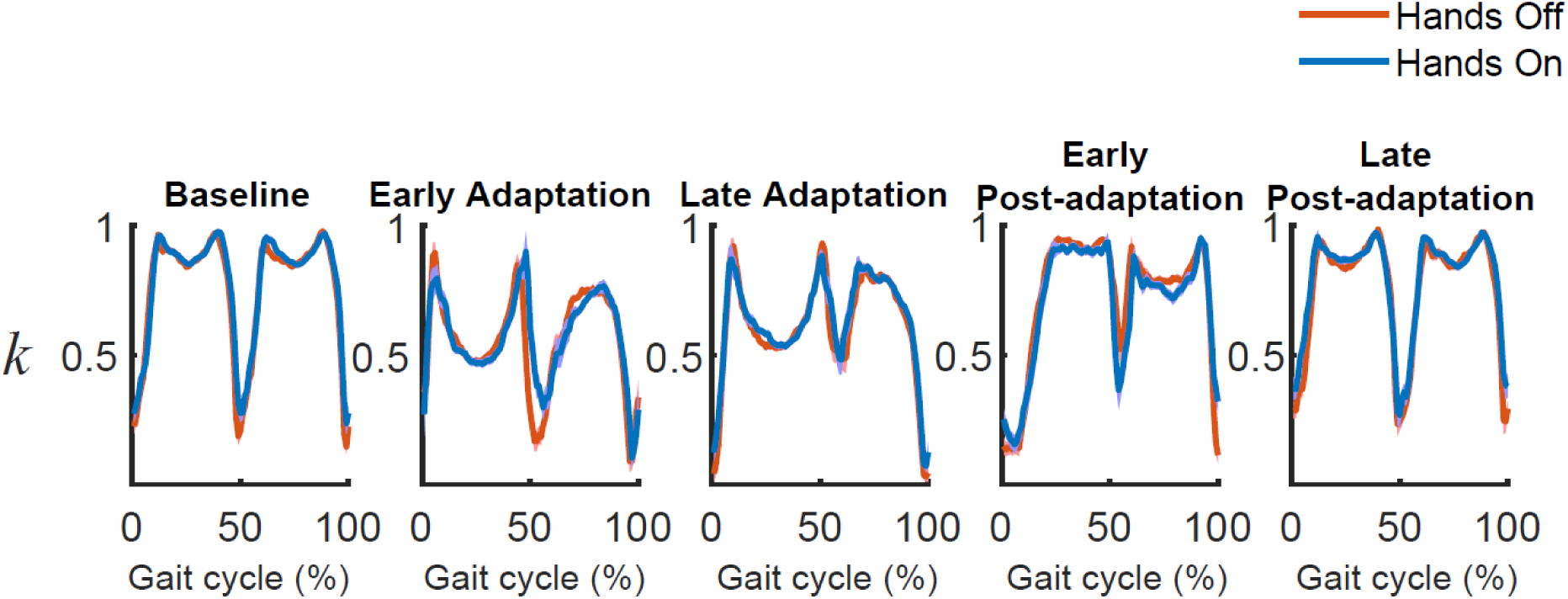
Limb cancellation (k) over the gait cycle during Baseline at 1.0 ms^-1^, Early Adaptation, Late Adaptation, Early Post-Adaptation, and Late Post-Adaptation. During Early and Late Adaptation, gait cycles start with foot strike on the slow belt such that the first half of the gait cycle corresponds to stance phase on the slow belt and the second half corresponds to stance phase on the fast belt. During Post-adaptation, the gait cycle begins with the step on the treadmill belt that was slower during Adaptation. The solid lines represent the median across participants and the shaded area around each curve represent the standard deviation across participants. Red: Hands Off. Blue: Hands On.

### Associations between SLA and dynamic balance

Measures of SLA were strongly associated with measures of WBAM during adaptation and post-adaptation. A two-way ANOVA revealed that there were significant main effects of Group (F = 3.07, p = 0.011) and Phase (F = 20.78, p < 0.001) on the correlation between SLA and *ΣL*_*fast*_, but there was no significant interaction between these factors (F = 0.59, p = 0.816). SLA was positively correlated with *ΣL*_*fast*_ during adaptation (Fig. 7A) and post-adaptation (Fig. 7B) for both the Hands Off and Hands On groups. Since the reduction in *ΣL*_*fast*_ during adaptation was represented as a positive change similar to SLA, the positive association between SLA and *ΣL*_*fast*_ indicates that momentum was reduced as asymmetry was reduced. The correlation between SLA and *ΣL*_*fast*_ was greater in the Hands Off group than the Hands On group (*r*_*Hands off*_ = 0.58 ± 0.29 versus *r*_*Hands On*_= 0.55 ± 0.23; p = 0.024; p = 0.024). The correlation was also greater during Post-Adaptation than Adaptation (*r*_*Adaptation*_ = 0.44 ± 0.25 versus *r*_*Post–Adaptation*_ = 0.69 ± 0.21; p = 0.001).

**Figure 7.**
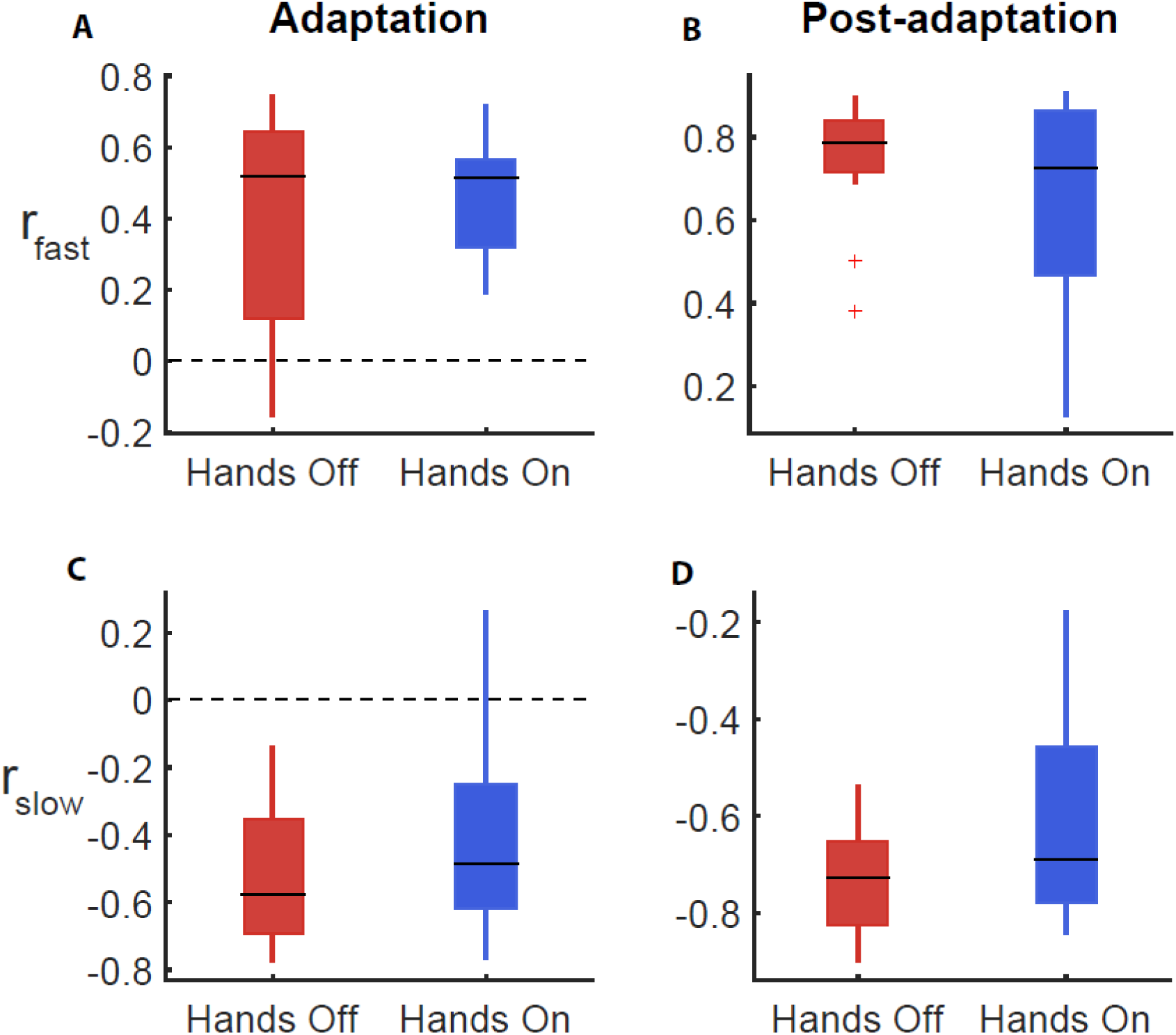
Associations between step length asymmetry (SLA) and measures of whole-body angular momentum during adaptation and post-adaptation. A. Correlations between SLA and integrated WBAM during the fast step during adaptation. B. Correlations between SLA and integrated WBAM over the fast belt during post-adaptation. C. Correlations between SLA and integrated WBAM over a step on slow belt for both groups during adaptation. D. Correlations between SLA and integrated WBAM over a step on the side which was slower during adaptation.

We also found main effects of Group (F = 5.09, p < 0.001) and Phase (F = 18.04, p < 0.001) on the correlation between SLA and *ΣL*_*slow*_, but there was no interaction between these factors (F = 0.6, p = 0.812). SLA was negatively correlated with *ΣL*_*slow*_ during adaptation (Fig. 7C) and positively correlated with *ΣL*_*slow*_ during post-adaptation (Fig. 7D). Each of these relationships indicate that momentum was reduced as asymmetry was reduced. Overall, the magnitude of the correlation between SLA and *ΣL*_*slow*_ was greater in the Hands Off group than the Hands On group (*r*_*Hands off*_ = -0.63 ± 0.19 versus *r*_*Hands On*_= -0.51 ± 0.28; p < 0.001). The correlation between SLA and *ΣL*_*slow*_ was greater during Post-Adaptation than Adaptation (*r*_*Adaptation*_ = -0.47 ± 0.25 versus *r*_*Post*-*Adaptation*_= -0.67 ± 0.19; p = 0.012). These results demonstrate that the provision of manual support leads to increased independence between asymmetry and stability.

## Discussion

In this study, we sought to determine the role of stability in driving adaptation to walking on a split-belt treadmill. We assessed stability using measures of whole-body angular momentum and found that the magnitude of angular momentum initially increased when people were exposed to the asymmetry in belt speeds but was rapidly reduced as people adopted more symmetric step lengths. In a second experiment, we provided participants with an external source of stability to determine how the addition of external stabilization influenced the adaptation process. When participants were held on to a handrail, we observed a reduction in step length asymmetry during Early Adaptation and during Early Post-Adaptation, and we also observed a weaker association between angular momentum and step length asymmetry. Together, these results suggest that the imposed asymmetry in belt speeds during split-belt adaptation acts as a perturbation to stability leading to increases in whole-body rotation. Moreover, the early reductions in asymmetry during the adaptation process may result, in part, from a process that seeks to stabilize whole-body dynamics.

### Regulation of whole-body momentum during split-belt adaptation

We observed systematic changes in whole-body angular momentum in response to the asymmetry conditions imposed by a split-belt treadmill. These changes in momentum resulted from perturbations of whole-body rotation and from changes in interlimb cancellation. During Adaptation, we observed a negative integrated momentum during steps on the fast belt and positive integrated momentum on the slow belt. Stepping on the fast belt, which moved more quickly than participants expected, resulting in an increase in forward pitch. People then recovered from this increase in forward pitch by rotating backward when stepping on the slow belt. When the belts were set to move at the same speed during post-adaptation, there was an after-effect in angular momentum that was opposite of what was observed during Early Adaptation. Specifically, during Early Post-Adaptation, integrated momentum was positive during steps on the belt that was faster during adaptation and was negative during steps on the belt that was slower during adaptation. Although the belts were set to move at the same speeds during post-adaptation, the belt that was moving slower during adaptation was now moving more quickly than expected and likely acted as a perturbation that resulted in an increase in forward pitch. This increase in pitch was offset by a positive angular momentum on the formerly fast belt to restore whole-body stability. Ultimately, both step length asymmetry and angular momentum returned to baseline levels by the end of the post-adaptation period.

The observed change in whole-body angular momentum during adaptation also resulted, in part, from step to step changes in the extent to which individual limb angular momenta offset one other. During Early Adaptation there was a reduction in the cancellation of angular momenta between the left and right sides and this contributed to the overall increase in whole-body angular momentum. As step length asymmetry was reduced during adaptation, participants reduced the magnitude of integrated angular momentum on each step by increasing limb cancellation. Although people achieved symmetric step lengths, the coefficient of limb cancellation was still smaller during Late Adaptation than Baseline. This suggests that a left-right asymmetry in segmental angular momentum remained due to the presence of left-right asymmetries in speed. After the asymmetry in belt speeds was removed, people reduced the magnitude of the integrated angular momentum close to zero as step length asymmetry decreased to zero and limb cancellation returned to the baseline level.

Our findings were consistent with previous studies looking at the whole-body angular momentum in the sagittal plane with people who naturally show an asymmetric walking pattern. For example, people post-stroke (Finley et al., 2015; Lauzière et al., 2014; Olney and Richards, 1996) or with unilateral transtibial amputees (Adamczyk and Kuo, 2015; Hak et al., 2014) often have a spatiotemporally asymmetric gait. In addition, whole-body angular momentum in the sagittal plane is greater than in age-matched controls in both of these groups (D’Andrea et al., 2014a; Honda et al., 2019; Pickle et al., 2014). These results, along with our findings, suggest that people who walk asymmetrically could have difficulty regulating angular momentum.

### The role of dynamic balance control during split-belt adaptation

Given our results, the role of balance control during locomotor adaptation is likely most evident during the initial adjustment to changes in belt speeds. We found that step length asymmetry and angular momentum were strongly associated with one another during the early phases of Adaptation and Post-adaptation. However, when participants held on to the handrail, the initial perturbation to step length asymmetry was reduced and the association between changes in step length asymmetry and changes in momentum was also weakened. Despite these changes during Early Adaptation, there were no changes in the final asymmetry that participants adopted during Adaptation. Taken together with previous work showing that reductions in asymmetry during adaptation lead to a more economical gait (Finley et al., 2013a; Sanchez et al., 2018; Selgrade et al., 2017), this suggests that early changes in SLA may promote dynamic stability while slower timescale reductions in SLA may be driven by energy optimization. Our findings are supported by recent evidence that people selectively adapt gait variables that lead to an increase in stability before modifying those that minimize energy expenditure (Cajigas et al., 2017).

Contrary to our initial hypothesis, whole-body angular momentum did not differ between handrail conditions during Adaptation or Post-Adaptation. Whole-body angular momentum is affected by both changes in body pitch and by the degree of cancellation of angular momentum between segments. Our analysis of segmental angular momentum revealed that holding on to a handrail had no effects on body pitch or cancellation of momentum between segments. This explains why we did not observe differences in the whole-body angular momentum between handrail conditions.

Given that there was no effect of holding the handrail on WBAM, it is likely that the effects of holding on to a handrail in reducing step length asymmetry reflect the contribution of light touch in balance control. Previous work has shown that lightly touching a handrail leads to a reduction in the variance of both fore-aft and mediolateral CoM position during walking on a treadmill (Dickstein and Laufer, 2004). This effect may result from light touch cues providing additional cues about the spatial location and orientation of the body which helps people reduce body sway. This has been shown previously during upright stance (Jeka, 1997; Jeka and Lackner, 1994). In the current study, holding on to the handrail may have reduced the initial perturbation to the body’s pitch in response to the unexpected increase in the speed of the fast belt. This would require a smaller recovery step on the slow belt since the foot could be placed closer to the body. If step timing is similar between the Hands On and Hands Free groups, the subsequent step on the fast belt would be longer for the Hands On group because the trailing limb on the slow belt would be in a more extended position. Either or both changes could be responsible for the observed reduction in step length asymmetry during early adaptation in the Hands On group.

### Conclusion

In this study, we examined how dynamic balance was regulated during adaptation to walking on a split-belt treadmill to understand the role of balance control on locomotor adaptation. Measures of dynamic instability scaled with asymmetry and gradually decreased over the course of adaptation. The strength of the relationship between measures of instability and asymmetry decreased with the provision of an external source of stability. Together, these results suggest that balance control is an important factor influencing locomotor learning, particularly when initially adjusting to changes in the environment.

